# *De novo* protein fold families expand the designable ligand binding site space

**DOI:** 10.1101/2021.01.13.426598

**Authors:** Xingjie Pan, Tanja Kortemme

## Abstract

A major challenge in designing proteins *de novo* to bind user-defined ligands with high specificity and affinity is finding backbones structures that can accommodate a desired binding site geometry with high precision. Recent advances in methods to generate protein fold families *de novo* have expanded the space of accessible protein structures, but it is not clear to what extend *de novo* proteins with diverse geometries also expand the space of designable ligand binding functions. We constructed a library of 25,806 high-quality ligand binding sites and developed a fast protocol to place (“match”) these binding sites into both naturally occurring and *de novo* protein families with two fold topologies: Rossman and NTF2. 5,896 and 7,475 binding sites could be matched to the Rossmann and NTF2 fold families, respectively. *De novo* designed Rossman and NTF2 protein families can support 1,791 and 678 binding sites that cannot be matched to naturally existing structures with the same topologies, respectively. While the number of protein residues in ligand binding sites is the major determinant of matching success, ligand size and primary sequence separation of binding site residues also play important roles. The number of matched binding sites are power law functions of the number of members in a fold family. Our results suggest that *de novo* sampling of geometric variations on diverse fold topologies can significantly expand the space of designable ligand binding sites for a wealth of possible new protein functions.

**Author summary:** *De novo* design of proteins that can bind to novel and highly diverse user-defined small molecule ligands could have broad biomedical and synthetic biology applications. Because ligand binding site geometries need to be accommodated by protein backbone scaffolds at high accuracy, the diversity of scaffolds is a major limitation for designing new ligand binding functions. Advances in computational protein structure design methods have significantly increased the number of accessible stable scaffold structures. Understanding how many new ligand binding sites can be accommodated by the *de novo* scaffolds is important for designing novel ligand binding proteins. To answer this question, we constructed a large library of ligand binding sites from the Protein Data Bank (PDB). We tested the number of ligand binding sites that can be accommodated by *de novo* scaffolds and naturally existing scaffolds with same fold topologies. The results showed that *de novo* scaffolds significantly expanded the ligand binding space of their respective fold topologies. We also identified factors that affect difficulties of binding site accommodation, as well as the relationship between the number of scaffolds and the accessible ligand binding site space. We believe our findings will benefit future method development and applications of ligand binding protein design.

## Introduction

Ligand binding is a major class of protein functions, and the ability to design ligand binding *de novo* has many important applications(1) such as engineering of biosensors and ligand-controlled protein functions(2, 3). Naturally occurring proteins recognize their cognate ligands with high affinity and specificity using defined three-dimensional geometries of binding sites with high shape complementarity between ligands and proteins. For the formation of favorable hydrophobic and polar interactions, the chemical groups on the protein must be placed at specific spatial positions relative to the ligand(4, 5). Designing new ligand binding proteins therefore requires the ability to build binding sites with defined geometries into stable protein scaffolds that can accommodate the desired interaction geometry with high accuracy. While this approach has led to the successful design of enzymatic activity(6, 7), ligand binding proteins(8, 9), and biosensors(2, 3, 10), it has been limited by both the availability of defined binding site geometries and stable protein scaffolds into which these binding sites can be placed(3).

Several methods have recently been developed to address the first problem, increasing the number of potential ligand binding sites one could generate. The RIF docking method(11) generates ensembles of billions of side chain placements that make defined hydrogen-bonding and non-polar interactions with a target ligand. Other methods(12, 13) use statistics from the protein data bank (PDB) to find three-dimensional placements of amino acid residues that form favorable interactions with fragments of a ligand, which can then be assembled into complete binding site geometries. Protein-ligand interactions defined by these methods have been built successfully into a *de novo* designed beta barrel(11), and a parametrically designed helical bundle(13).

Naturally occurring proteins solve the second problem, finding a suitable protein backbone to accommodate a specific binding site geometry, not by using a different fold for each function but instead by evolving structural variation in existing protein fold families. This variation allows proteins with the same fold topology (identity and connectivity of secondary structure elements) to tune the precise geometry of binding sites to recognize diverse ligands(14). This strategy has recently been mimicked by advances in computational protein design methods. These methods have generated *de novo* designed protein fold families with large numbers of diverse geometries(15, 16), which have significantly expanded the accessible designable protein structure space. The resulting *de novo* proteins might be able to support binding sites that cannot be built onto naturally occurring proteins in the PDB, but the extent to which *de novo* fold families could improve binding site design has not been explored. Understanding the relationship between the space occupied by protein structures, and the space available to support different functions, is important for developing methods to design proteins *de novo* that can bind to novel and highly diverse user-defined ligands.

Here, we studied the ability of native and *de novo* fold families to support a large number of different ligand binding sites. We built a high-quality ligand binding site library from high resolution protein crystal structures. We then matched the binding site library to members of protein folding families using two protocols: a newly developed “fast matching” protocol and the standard method for matching in the Rosetta program for structure modeling and design(5). We calculated the number of matched binding sites for four fold families with two different topologies. We studied the effects of binding site sizes, ligand sizes and primary sequence separation of binding site residues on the matching success rates and determined the increase of numbers of matches with increasing the sizes of fold families. Together, we show that *de novo* fold family design is a promising approach to broaden the scope of designable ligand binding sites.

## Results

We first constructed a library of ligand binding sites from native proteins in the PDB. We extracted 25,806 ligands that have at most 100 heavy atoms as well as the ligand binding site residues from 23,238 cluster representative structures from the PDB95 database(17) where chains from the protein data bank are clustered at 95% identity (**Methods**). The extracted ligands have between 1 and 93 heavy atoms (**Fig 1A,B**). 80.6% percent of the ligands have 13 or fewer heavy atoms, and 7,335 (28.4%) of the ligands have only 1 heavy atom. There are 2,461 unique ligand types in the 25,806 binding sites. The distribution of ligand type frequencies has a long tail (**S1 Table**). There are 33 frequent ligand types that appear in over a hundred binding sites, while 1,817 ligand types only appear in single binding sites. The frequent ligand types include common crystallographic additives such as glycerol; 1,2-ethanediol; ions such as SO_4_^2−^ and Mg^2+^; and cofactors such as heme and flavin adenine dinucleotide (FAD). Ligands that appear in multiple binding sites are seen as vertical stripes in **Fig 1B**. Binding sites have between 2 and 41 residues, with 81.2% of the binding sites having 7 or fewer binding site residues. The number of protein residues in binding sites scales linearly with the number of ligand heavy atoms, with a slope 0.35 (**Fig 1B**). The frequencies of amino acid types in binding sites are different from those for whole proteins reported by UniProtKB/Swiss-Prot (**Fig 1C**). We defined the enrichment ratios of amino acids as their frequencies in ligand binding sites divided by their frequencies in whole proteins. The large aromatic side chains Trp, Tyr and Phe are the top 1, top 3 and top 6 enriched amino acid residues, respectively. His, characterized by its ability to coordinate metal ions and to catalyze chemical reactions, is the second most enriched amino acid residue. Asp and Arg are the 4th and 5th enriched amino acid residues, which may play important roles in interacting with charged ligands. Binding sites with single heavy atom ligands have different amino acid preference than those binding to ligands with at least two heavy atoms (**Fig 1D**). For the binding sites with single heavy atom ligands, the negatively charged residues Asp and Glu, which can form favorable electrostatic interactions with positively charged metal ions, are highly enriched. The enrichment ratios of Asp and Glu are 4.6 and 2.2, respectively. The top 5 enriched residues that bind to ligands with at least 2 heavy atoms are Trp, His, Tyr, Phe and Arg.

**Fig 1.**
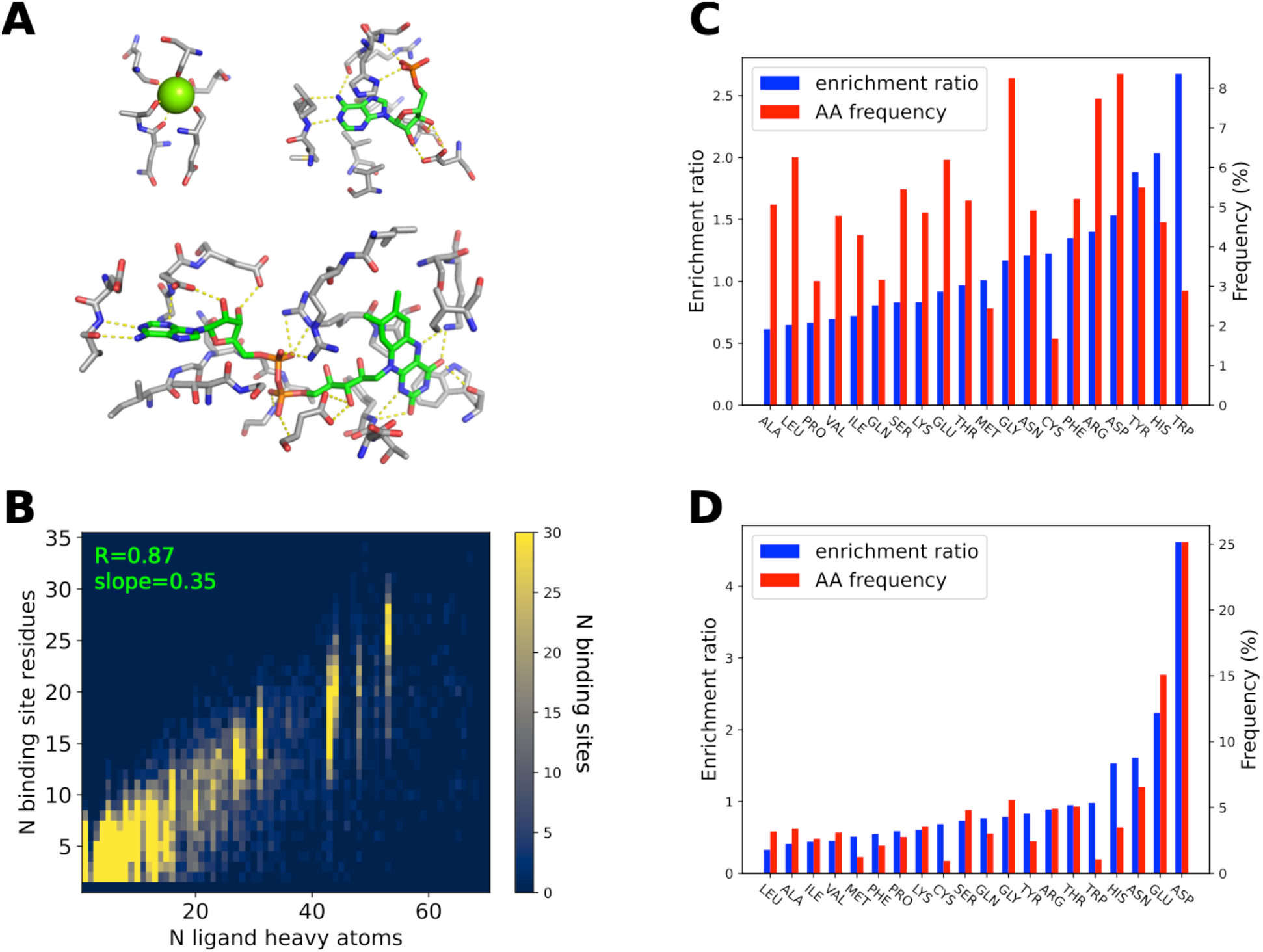
The ligand-binding site library. **A.** Binding site examples. The Mg^2+^ ion is shown as a sphere; small molecules and protein residues are shown as sticks; carbon atoms are colored in green (small molecule) or grey (protein residues); oxygen atoms are colored in red; nitrogen atoms are colored in blue; polar interactions are shown as yellow dashed lines. **B.** Joint distribution of binding site sizes (numbers of binding site protein residues) and numbers of ligand heavy atoms. Binding site sizes are linearly correlated with the numbers of ligand heavy atoms. **C, D.** Amino acid (AA) frequencies (red, right y-axis) in ligand-binding sites and enrichment ratios (blue, left Y-axis) in ligand-binding sites compared to all residues in a protein. **C.** Distributions of all ligand binding sites. **D.** Distributions of single heavy atom ligand binding sites.

The binding site library is useful for testing the ability of protein fold families to support ligand binding sites. A protein scaffold can in principle support a ligand binding site if the binding site residues can be built onto the scaffold such that the key interactions between the ligand and binding site protein residues are preserved. The Rosetta matcher protocol(5) has been shown to be successful in matching ligand binding sites to protein scaffolds(8). However, the Rosetta matcher is too slow to match tens of thousands of binding sites to thousands of scaffolds because it samples all possible side chain rotamers of binding site residues. To perform all-against-all matching between the library of ligand binding sites and the sets of scaffolds, we developed a new fast match protocol (**Fig 2A**). In the fast match protocol, the binding site is anchored and matched as a rigid body (**Methods**). This rigid body approximation drastically improved the matching speed. We tested the run time by matching the binding site library to the native NTF2 fold family (CATH superfamily 3.10.450.50)(18). The mean time to find a successful standard Rosetta match is 706 s while the mean time of a successful fast match is 3.1s. As a trade-off, the rigid body approximation of the fast match method may discard binding sites that can be matched by the Rosetta matcher using alternative side chain rotamers. Therefore, in this study we focused on matching ligand binding sites using the side chain rotamers present in the original ligand binding site in the PDB. Using these original rotamers also let us directly compare the backbone geometries in the native binding sites and the backbone geometries in our scaffold libraries.

**Fig 2.**
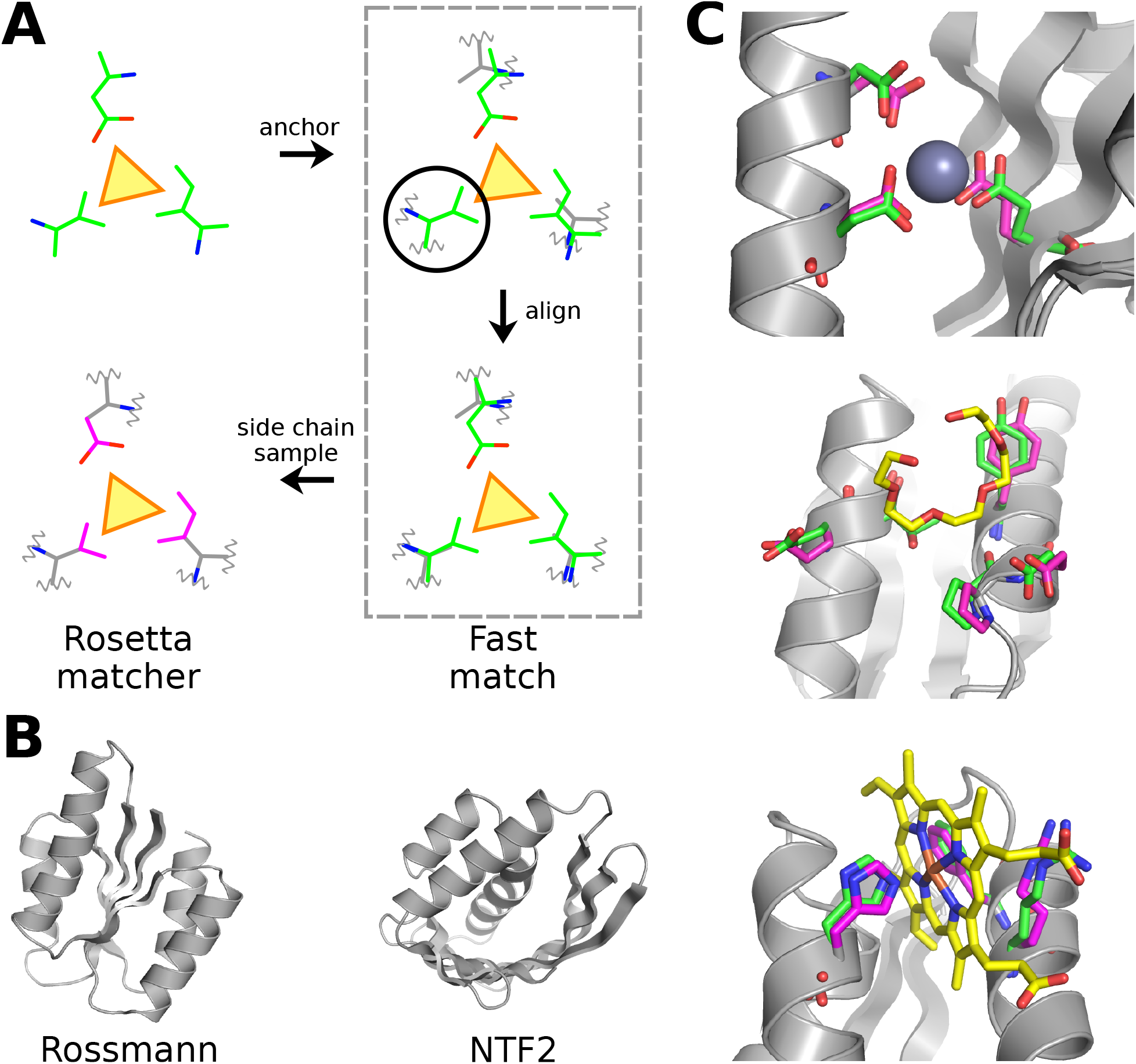
Matching ligand binding sites to scaffold libraries. **A.** Schematic of the matching protocol. The ligand is represented as a yellow triangle. The ligand-binding site as a rigid body (green) is first matched to the scaffold (grey) by anchoring to a scaffold residue shown in the black circle. Then the binding site residues are aligned to the corresponding scaffold residues. Finally, the standard Rosetta matcher is applied to build the binding site side chains (magenta) onto the scaffold. **B.** The binding sites are matched to native and *de novo* designed scaffold families with Rossmann or NTF2 fold topologies. **C.** Examples of matches. The coloring scheme is the same as **A**.

We matched the binding site library to backbone scaffolds of *de novo* designed Rossmann and NTF2 protein fold families generated by the loop-helix-loop unit combinatorial sampling (LUCS) method(15), as well as the two native fold families with the same topology from the CATH database(18) (**Fig 2B, Methods**). To determine if a fold family can support a given ligand binding site, we first used fast match to match the ligand binding site to all protein scaffolds in the family. Then we used the Rosetta matcher to match the binding site to the scaffolds that passed the fast match (**Methods**). To limit computational time, once the Rosetta matcher found a match for a given binding site, we skipped matching to further scaffolds in the same family. Since we used stringent matching criteria (**Methods**), the matched binding sites in the scaffold closely recapitulated the interactions between the ligands and binding site residues in the original protein structures from which the binding sites were derived (**Fig 2C**).

Between 5896 and 7548 binding sites could be successfully matched by the Rosetta matcher to each fold family when considering all binding sites (**Table 1**). The number of binding site residues was the major determinant of the matching success rate (**Fig 3**). For the *de novo* Rossmann fold family, the success rates for 2, 3 and 4 protein residue binding sites were 93.8%, 33.4% and 6.5%, respectively. Only 13 binding sites with 5 or 6 residues could be matched. There was no match for binding sites with more than 6 protein residues. Similar dependencies on binding site sizes were observed across the 4 different protein fold families (**Table 2**). Because almost all 2-residue binding sites could be matched and the matching success rates were low for binding sites with more than 3 residues, we used 3-residue binding sites to further study properties of successful matches. We constructed a new library of binding sites that all have 3 protein residues (**Methods**) and matched the binding sites to the scaffold libraries using the same protocol. The number of successfully matched 3-residue binding sites ranged from 2,142 to 3,715 (**Table 1**).

**Table 1.** Number of matched binding sites.

| Binding site library | Match type | Native Rossmann | Native NTF2 | De novo Rossmann | De novo NTF2 |
| --- | --- | --- | --- | --- | --- |
| All binding sites | fast | 6860 (248)* | 8761 (795) | 9034 [2442]** | 8909 [943] |
|  | Rosetta | 5896 (212) | 7450 (580) | 7475 [1791] | 7548 [678] |
| 3 protein residue binding sites | fast | 3556 (324) | 5714 (1306) | 6537 [3305] | 6128 [1720] |
|  | Rosetta | 2142 (199) | 3541 (807) | 3715 [1772] | 3686 [952] |

\* Numbers in parentheses are binding sites that cannot be matched to *de novo* scaffolds with the same topology.
\*\* Numbers in square brackets are binding sites that cannot be matched to native scaffolds with the same topology.

**Table 2.** Dependency of matching success on binding site size (number of protein residues)

| Binding site size | Native Rossmann |  | Native NTF2 |  | De novo Rossmann |  | De novo NTF2 |  |
| --- | --- | --- | --- | --- | --- | --- | --- | --- |
|  | success count | success rate | success count | success rate | success count | success rate | success count | success rate |
| 2 | 4590 | 80.9% | 5340 | 94.2% | 5328 | 93.8% | 5359 | 94.4% |
| 3 | 1182 | 21.4% | 1792 | 32.5% | 1853 | 33.4% | 1882 | 33.9% |
| 4 | 118 | 2.7% | 272 | 6.3% | 281 | 6.5% | 276 | 6.4% |
| 5 | 6 | 0.2% | 38 | 1.4% | 12 | 0.4% | 27 | 1.0% |
| 6 | 0 | 0 | 6 | 0.4% | 1 | 0.06% | 3 | 0.2% |
| 7 | 0 | 0 | 2 | 0.2% | 0 | 0 | 1 | 0.1% |

**Fig 3.**
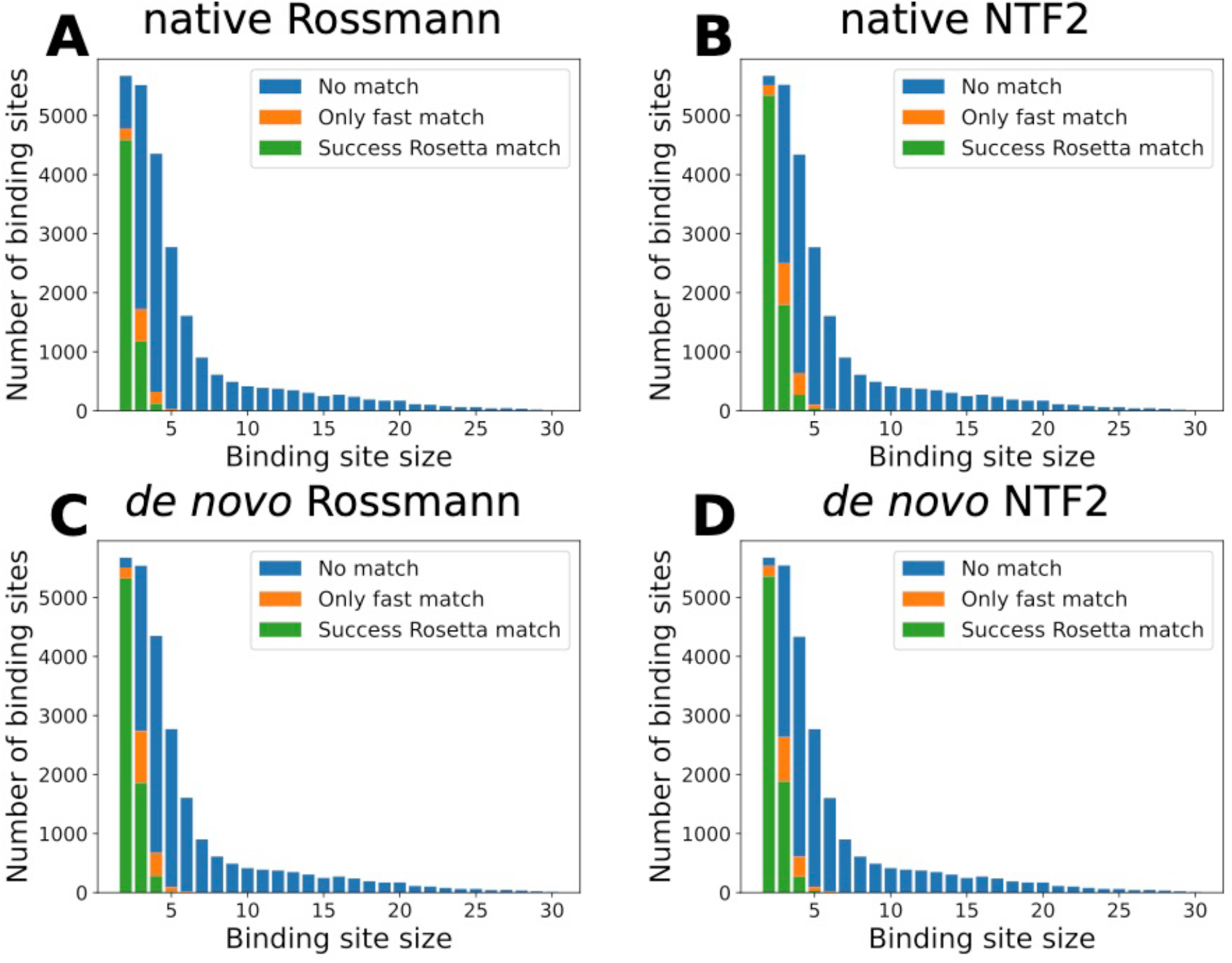
Matchability of ligand binding sites depends on the binding site size. Histograms of numbers of matches vs binding site sizes (number of protein residues in the binding site). Bindings sites that cannot be matched to any scaffold are shown in blue. Bindings sites that can be matched to at least one scaffold by the fast match method but cannot be matched by the standard Rosetta matcher are shown in orange. Binding sites that can be matched to at least one scaffold by the standard Rosetta matcher are in green. **A-D.** Results for 4 scaffold libraries; scaffold sets are indicated in each panel title.

For the successfully matched 3-residue binding sites, we first analyzed the positions of matches relative to the surface of the scaffolds. For each scaffold, we used the Rosetta Layer residue selector(19) to assign layers to all of its residues in a side chain independent manner (**Methods**). Residues on the surfaces of scaffolds were assigned to the surface layer; deeply buried residues were assigned to the core layer; and the rest of residues were assigned to the boundary layer (**Fig 4A**). In all of the fold families, surface layer residues were most abundant, which accounted for 47%-63% of all residues. 29%-39% residues were in the boundary layer and 6%-20% residues were in the core layer (**Fig 4B**). NTF2 fold proteins had more surface layer residues which was likely due to the pocket of this fold. We defined the layer of each residue in a matched binding site as the layer of its matched scaffold residue position. The frequencies of matched residue layers are similar to the frequencies of scaffold layers (**Fig 4C**). To evaluate the positions of matches at the binding site level, we defined a depth score for each matched binding site. The depth score of a matched binding site is the number of boundary residues plus two times the number of core residues. The depth scores for binding sites matched to different fold families had similar distributions (**Fig 4D**). 20%-27% binding sites were entirely matched to protein surfaces and had depth scores of 0. The remainder of matched binding sites were buried to some extent. The majority of binding sites were in shallow pockets with depth scores ranging from 2 to 4. Only 8%-12% binding sites were matched to deeply buried positions with depth scores of 5 or 6.

**Fig 4.**
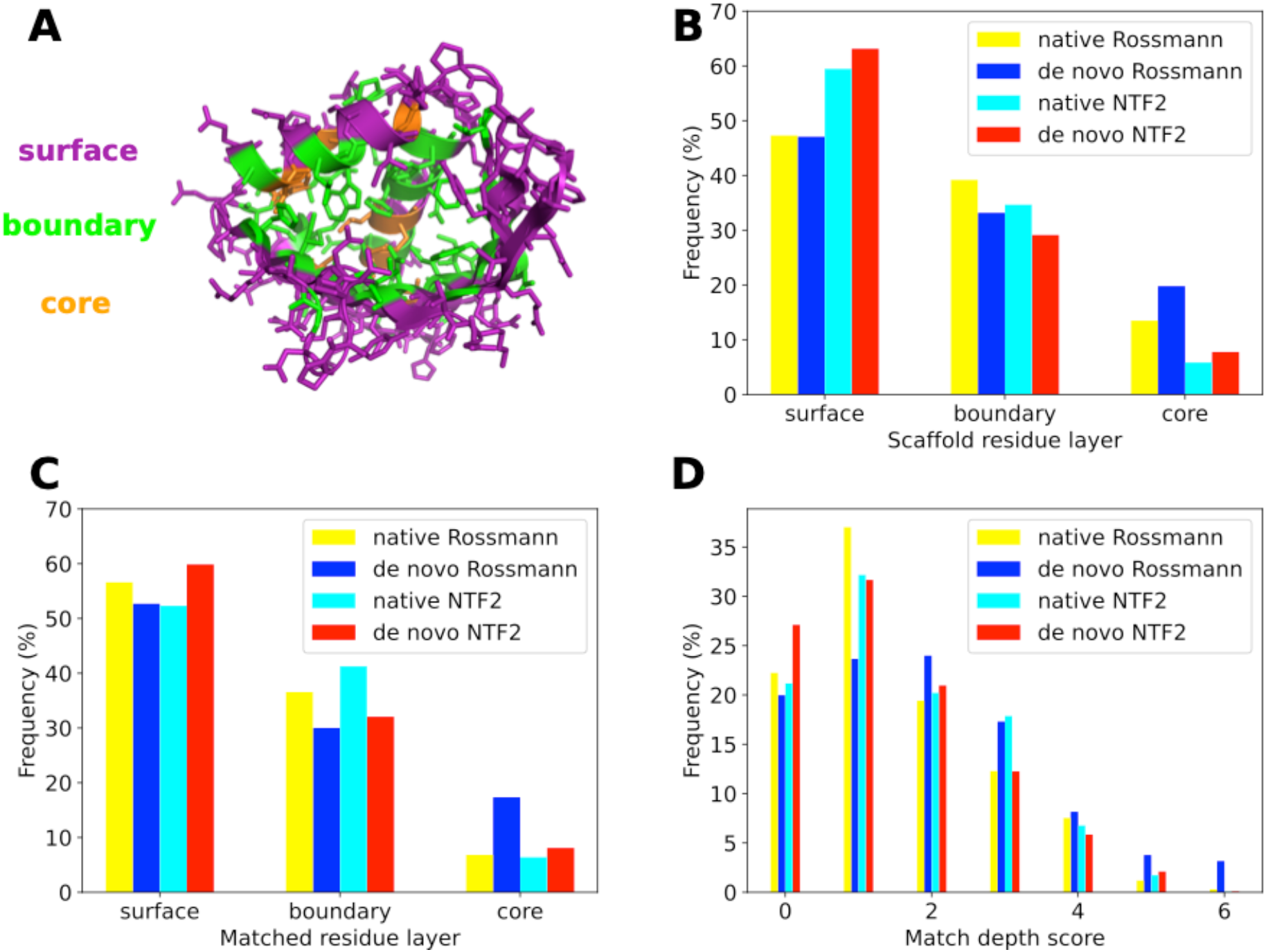
Ligand binding sites are matched to all layers of scaffolds. **A.** An example of scaffold residue layers assigned to a scaffold (PDB:3FH1) from the native NTF2 fold family by the Rosetta Layer residue selector. The surface, boundary and core layers are colored in purple, green and orange, respectively. **B.** Distributions of residue layers in different scaffold libraries. **C.** Distributions of residue layers of binding sites matched to different scaffold libraries. **D.** Distributions of binding site depth scores matched to different scaffold libraries.

We then tested factors that affect the “matchability” of 3-residue binding sites. We compared the number of overlapping binding sites that were matched to both of two fold families to the expected number of overlapping binding sites (**Fig 5A**). If matching to one fold family is independent from matching to another fold family, the probability of overlapping binding sites should be the product of the probabilities of matching to each fold family. We compared 4 pairs of scaffold libraries (**Fig 5A**): *de novo* designed Rossmann folds versus *de novo* designed NTF2 folds (top left) or versus native Rossmann folds(top right), and native NTF2 folds versus *de novo* designed NTF2 folds (bottom left) or versus native Rossmann folds (bottom right). For all 4 pairs of scaffold libraries, the observed number of overlapping binding sites was significantly higher than the number of expected overlapping binding sites (chi-squared test p-value < 10^−300^). This result indicates that some binding sites had higher matchabilities (probabilities to be matched to multiple scaffold libraries). We investigated the contribution of ligand sizes to binding site matchabilities. As expected, the matching success rates for 3-residue binding sites decreased with an increase of the number of ligand heavy atoms (**Fig 5B**), likely because larger ligands are more likely to clash with the scaffold backbones. We also hypothesized that binding sites whose residues have larger separations in primary sequences are more difficult to match. To confirm that non-local binding sites are harder to match, we calculated the mean inter-residue primary sequence distances for each 3-residue binding site and plotted the mean distances against the matching success rates (**Fig 5C**). When the 3-residues in a binding site were consecutive in primary sequence, the mean primary sequence distance was 1.33, and the matching success rates were higher than 80%. The success rates dropped rapidly with the increase of mean distance and reached a plateau at low match success rates when the mean distance reached 70.

**Fig 5.**
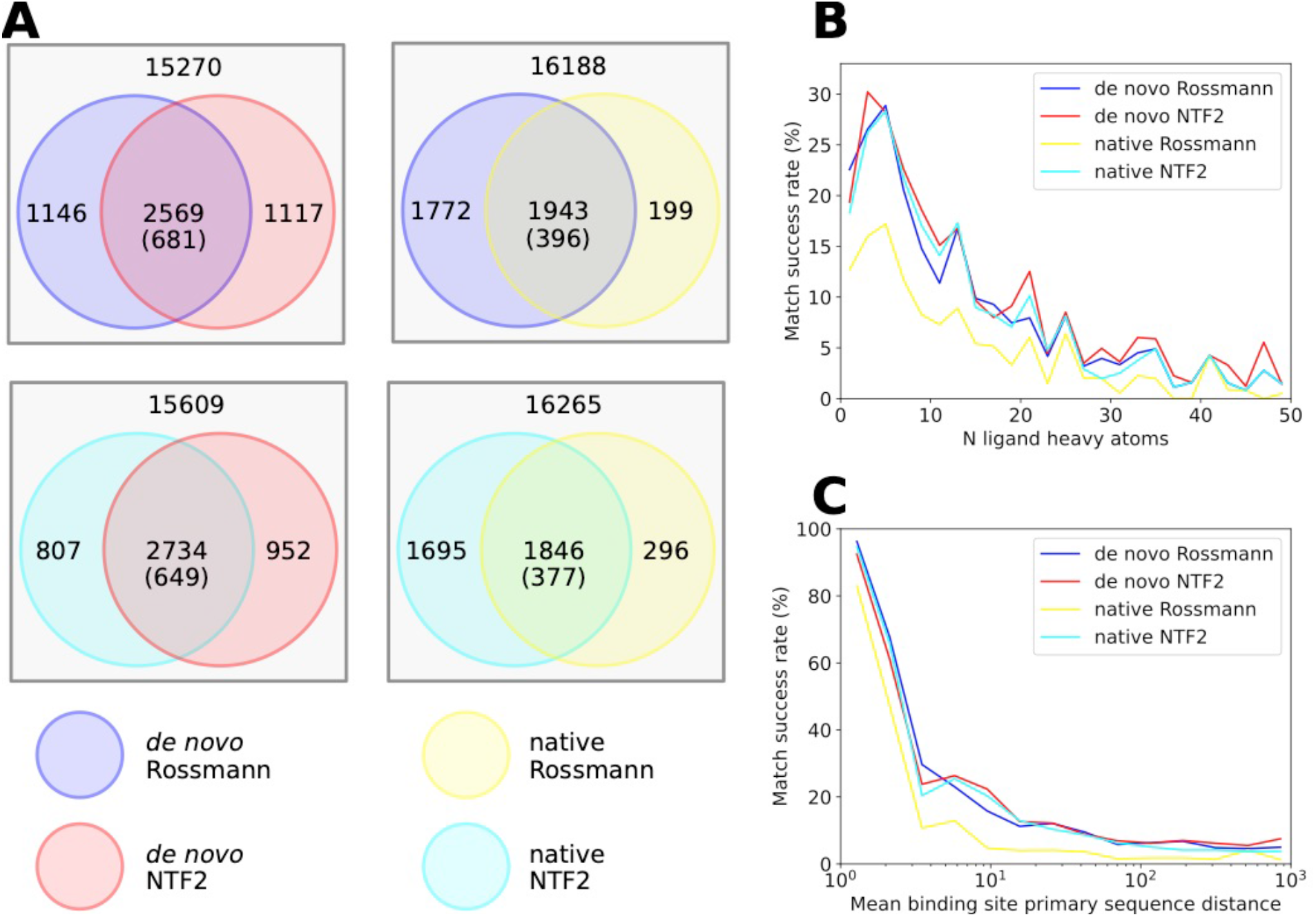
Features affecting matching success rates of 3-residue ligand binding sites. **A.** Venn diagrams of the number of Rosetta-matched 3-residue binding sites between pairs of scaffold sets. The number in the overlapping region is the observed number of binding sites that can be matched to both scaffold sets, with the expected number in parentheses. The number in the non-overlapping region within a circle denotes the binding sites that can only be matched to this scaffold set. The number outside the circles denotes the binding sites that cannot be matched to either of the two scaffold sets. **B.** The numbers of ligand heavy atoms are negatively correlated with the match success rates. **C.** The mean primary sequence distances between binding site residues are negatively correlated with match success rates.

Next, we studied how the number of matched 3-residue binding sites grew with an increase of the number of scaffolds in fold families (**Methods**). The log of the number of matched binding sites scaled linearly with the log of the number of scaffolds (**Fig 6A-D**). This power law relationship was valid for both the number of fast matches and Rosetta matches across the 4 different fold families. The powers of the power law functions (slopes of the log-log plots) ranged from 0.184 to 0.298. Since the powers were small, the increase of matches progressively diminished as the number of scaffolds got large. Because there is a limited number of designable structures for each fold family, the power law relationship cannot continue indefinitely, but it can still provide a reasonable estimation of the upper bound of the number of matches. Extrapolating the *de novo* fold family power law relationships to the number of representative structures from the PDB95 database, i.e., 23,238 structures, the numbers of expected Rosetta matches for the Rossmann fold family and the NTF2 fold family would be 7,346 and 6,640. Based on this analysis, the extrapolated numbers of matchable binding sites are still much smaller than the number of total binding sites, highlighting the importance of having diverse fold topologies for different functions.

**Fig 6.**
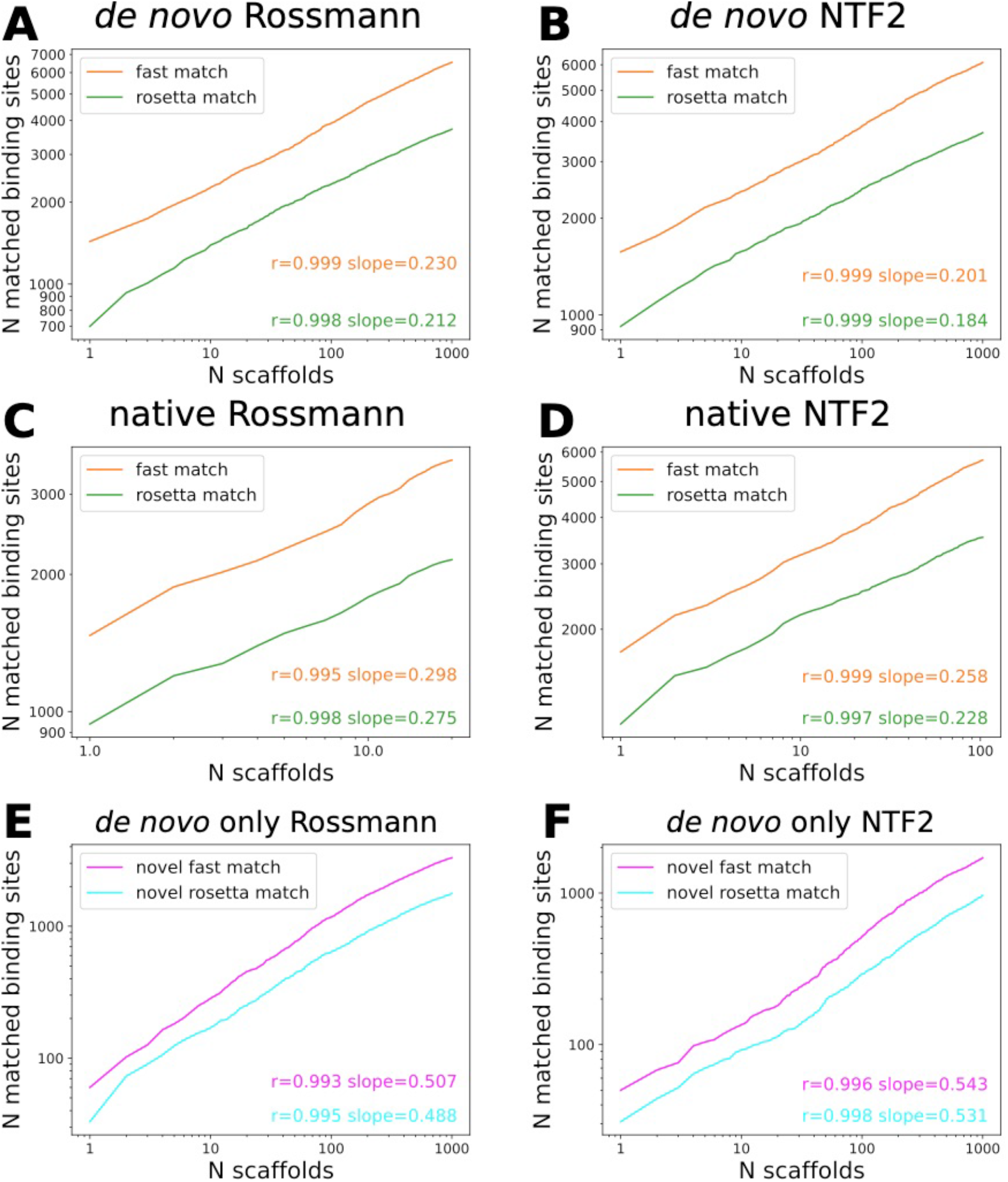
Numbers of matches scale as power-law functions of numbers of scaffolds in fold families. **A-D.** Log-log plots of the number of 3-residue matches vs the number of scaffolds. **E-F.** Log-log plots of the number of 3-residue binding sites that can only be matched to *de novo* scaffolds of specific topologies vs the number of scaffolds.

Finally, to understand how *de novo* scaffolds expand protein function space, we studied the binding sites that can be matched to *de novo* scaffolds but not to the native scaffolds of the same topology. We plotted the number of binding sites that were matched to only *de novo* fold families versus the number of *de novo* scaffolds (**Fig 6E,F**). For each topology, there are more than 1,000 binding sites that are exclusively matched to *de novo* scaffolds. These relationships also follow power law functions. The slopes are larger than the slopes of the total matches (**Fig 6A-D**), indicating that binding sites that can match to both native and *de novo* fold families saturate quickly.

## Discussion

Advances in computational protein structure sampling methods(20) have expanded the accessible structure space of *de novo* designed proteins. In particular, two recently developed computational methods(15, 16) are capable of engineering *de novo* protein families that contain defined variations in geometry of proteins that share the same overall fold topology. We probed the functional implications of *de novo* protein fold families generated by the LUCS method(15) by matching known ligand binding sites to both native and *de novo* fold families. We found that thousands of ligand binding sites that cannot be matched to native fold families can be matched to LUCS-generated members of *de novo* fold families of the same topology, showing that LUCS generated structures expand both the accessible protein structure and the accessible protein function space. The number of matched binding sites increased as a power law function of the number of scaffolds. This relationship allowed us to estimate the upper bound of matches as the number of scaffolds grew and showed that, in addition to geometric variation, different fold topologies are necessary to support diverse functions.

Previous studies have shown that computationally generated artificial (ART) compact homopolypeptide structures can match virtually every native ligand binding pocket(21, 22). In contrast, the native and *de novo* fold families we studied here can only be matched to a limited fraction of native binding sites. A likely reason is that we only used structures with two topologies while the ART structures are generated using secondary structure preferences from thousands of random PDB structures with many different fold topologies. These two behaviors together support that the diversity of topologies is important for the repertoire of native ligand binding functions. Additionally, the *de novo* designed structures we used were subjected to filters for a set of physical properties such as core packing, hydrogen bonding and surface exposed hydrophobic patches(15). These filters are designed to eliminate structures that are not likely to fold, whereas the ART structures are model polyleucine homopolypeptides. Requirements for folding places diverse additional constraints on the accessible conformational space of protein structures.

Using the new fast match protocol introduced here as well as the Rosetta matcher, we were able to match a library of high-quality binding sites to *de novo* protein fold families. To engineer new ligand binding proteins, the matching step is typically followed by sequence design(3, 8) to optimize the binding site protein environment. Ligand binding site design is a challenging problem because the designed sequence must simultaneously be compatible with the protein fold and precisely place binding site residues in their desired geometries for favorable interactions with the ligand. Given the typically high stability of *de novo* designed protein families(15, 23), matches generated by the protocol described here could be good model systems for testing binding site design algorithms.

Another advantage of using *de novo* fold families for ligand binding site design is that the systematic sampling of diverse geometries could provide an ensemble of negative states. Using negative states in design has been shown to improve accuracy in protein stability prediction(24). Thus, a *de novo* ensemble of negative states may increase success rates of ligand design where high accuracy in both sampling and scoring designs is required. Ensembles of different conformational states in *de novo* fold families also pave the way to engineer ligand binding-induced conformational changes. Small molecule-induced switches could be designed by building a ligand binding site in one of the structures in the *de novo* fold family and tuning the free energy gaps between the ligand binding state and the other states. We envision that *de novo* designed protein fold families will play an important role in designing functions such as ligand binding and protein switches.

## Methods

### Binding site library construction

Ligand binding sites were extracted from the PDB95(17) database. The representative pdb structures for each cluster, which were listed in the pdb_95.cod file, were used for binding site extraction. The representative structures were filtered by resolution. Only crystal structures whose resolutions were better than 2 Å were kept. Ligand residues were identified by built-in functions in PyRosetta(25). In this study, we focused on ligands that had at most 100 heavy atoms. Ligands that had average heavy atom B-factors greater than 60 Å^2^ were filtered out. Ligands that did not have protein residues within 5 Å were also excluded from subsequent processing. We calculated the Rosetta 2-body energy scores(26, 27) between ligands and protein residues that have at least one heavy atom within 5 Å from any ligand heavy atom. Ligand binding site residues were defined as protein residues that had favorable van der Waals, electrostatic or hydrogen bond interactions with the ligand. A residue was included in a binding site if the sum of its Rosetta energy(27) terms fa_atr, fa_elec, hbond_bb_sc and hbond_sc was less than −1 Rosetta energy units (REU). We excluded protein residues from consideration that had total Rosetta scores greater than 50 to avoid poorly modeled residues, such as those who have severe clashes with the protein environment. We also excluded all residues with missing heavy atoms in the PDB file. We only kept ligand binding sites that have at least two protein residues. To prevent overcounting ligands in structures which had multiple chains of the same protein in their asymmetric units, only one binding site was extracted for the same ligand in a given structure.

### Fast match protocol

We developed a new fast match protocol to rapidly match the library of binding sites to the sets of protein scaffolds. During the fast match, a ligand binding site is treated as a rigid body. When the fast matcher matches a ligand binding site to a scaffold, it first iterates through all pairs of binding site protein residues and scaffold residues. For each pair of residues, the protocol superimposes the N, Ca and C atoms of the binding site residue to the corresponding atoms in the scaffold residue. The remainder of the binding site is transformed as a rigid body. Then the matcher finds the closest scaffold residues to each binding site protein residue. The distances between residues are defined as the Ca-Ca distances. If all distances between binding site protein residues and their closest scaffold residues are within 2 Å, the backbone N, Ca and C atoms of the binding site protein residues are superimposed to the N, Ca and C atoms of their closest scaffold residues. The superimposition minimizes the root mean squared deviation (RMSD) between the corresponding atoms. If the RMSD is within 1 Å, the cosine of angles between the vectors pointing from Ca to Cb of corresponding residues are calculated. If all the cosine values are greater than 0.7, clashes between the matched binding site and the scaffold backbone are checked. Two atoms are defined to clash when the distance between them is less than the sum of their Lennard-Jones radii times a scale factor of 0.6. The match is accepted if the ligand and protein side chains from the binding site do not clash with the scaffold backbone atoms that are not matched to binding site residues.

### Standard Rosetta matcher

For each binding site successfully matched to a scaffold using fast match, we ran the standard Rosetta matcher(5). We made mol2 files for ligands using Open Babel(28) and generated ligand parameter files with the molfile_to_params.py script distributed with Rosetta. The relative positions of a ligand and a binding site protein residue are defined by 6 heavy atoms. On the ligand side, the heavy atom closest to the protein residue and the two ligand heavy atoms closest to the first ligand heavy atom are defined as the anchor atoms. On the protein residue side, the heavy atom closest to the ligand and two protein atoms closest to the first protein heavy atom are defined as the anchor atoms. For each binding site, we generated a constraint file where the relative positions between the ligand anchor atoms and protein residue anchor atoms were constrained. We used stringent matching criteria similar to those used in previous work(8, 12). The relative distances between ligands and binding site residues are sampled at ideal values; the relative angles and torsions are sampled at the ideal values and ±10° from the ideal values. The binding sites were matched using the standard Rosetta matcher with the following command:

match.linuxgccrelease -match:output_format PDB -match:match_grouper SameSequenceGrouper -match:consolidate_matches -match:output_matches_per_group 1 - use_input_sc -in:ignore_unrecognized_res -ex1 -ex2 -enumerate_ligand_rotamers false - match∷lig_name LIG_NAME -match:geometric_constraint_file CST_FILE -s SCAFFOLD_PDB - match∷scaffold_active_site_residues POS_FILE

where LIG_NAME is the 3-letter name of the ligand, CST_FILE is the constraint file, SCAFFOLD_PDB is the pdb file of the scaffold structure and POS_FILE is the file that stores the matchable residues. In this study, all residues on a scaffold are matchable.

### Construction of scaffold libraries

The *de novo* Rossmann and NTF2 fold families were reported in ref.(15). The scaffolds in these fold families were generated by the LUCS method and filtered by a set of designability filters(15). We randomly selected 1,000 scaffolds from each *de novo* fold family as the scaffold set for ligand binding site matching. The native fold families of Rossmann and NTF2 folds were obtained from the CATH database(18). The native Rossmann fold scaffolds were extracted from the CATH 3.40.50.1980 superfamily and the native NTF2 family structures were from the CATH 3.10.450.50 superfamily. Because the automatic classification algorithm of the CATH database did not force all structures in a CATH superfamily to have a same topology, we manually excluded the CATH structures that have different topologies from the *de novo* designed scaffolds. As a result, the native Rossmann fold scaffold set had 20 structures and the native NTF2 fold scaffold set had 103 structures. The C-terminal helices in *de novo* NTF2 scaffolds occluded the ligand binding pocket. In contrast, only 35 out of 103 native NTF2 scaffolds had C-terminal helices. Among these native C-terminal helices, 31 helices pointed away from pocket entrances, and thus, did not affect the accessibility of ligand binding sites, leaving only 4 scaffolds with pocket occluding C-terminal helices. We therefore trimmed the C-terminal helices in *de novo* NTF2 proteins to expose the ligand binding pocket.

### Construction of a library of 3-residue binding sites

The 3-residue binding site library was constructed from the library of all binding sites. We eliminated binding sites with fewer than 3 residues. The binding sites with 3 protein residues were kept unchanged. For binding sites with more than 3 protein residues, we scored the total Rosetta two-body energy(26) between the ligand and each protein residue. We kept the 3 protein residues with lowest total two-body energies and removed the remainder of the binding site residues.

### Assignment of layers to scaffold residues

The Rosetta Layer selector(19) with the default settings was applied to assign layers to each scaffold residue. The layer of a residue was determined by a weighted count of the number of neighbor amino acid residues in a cone extending along its Ca-Cb vector. A residue is assigned to the surface layer if the weighted count is less than 2; a residue is assigned to the core layer if the weighted count is greater than 5.2; all other residues are assigned to the boundary layer.

### Calculation of the numbers of matches for subsets of fold families

During the process of matching a binding site to a fold family, we recorded the number of scaffolds in the fold family that we tested to find the first successful fast match and called this number the first-fast-match-encounter-number. The number of fast matches for a subset of a fold family with N scaffolds was defined as the number of binding sites with first-fast-match-encounter-numbers smaller than or equal to N. The number of Rosetta matches for subsets of fold families were calculated in the same way.

## Supporting information

S1 File. Summary tables of matching results to all fold families.

## Data availability

All relevant data are available in the manuscript and supporting information data files. Rosetta source code is available from rosettacommons.org. Scripts, the binding site library and the scaffold sets are available at https://github.com/Kortemme-Lab/match_ligand_binding_sites/releases/tag/v1.

## Acknowledgments

This work was supported by grants from the National Institutes of Health (NIH) (R01-GM110089) and the National Science Foundation (NSF) (DBI-1564692) to TK. We additionally acknowledge a UCSF Discovery Fellowship to XP. TK is a Chan Zuckerberg Biohub Investigator.

## Author contributions

XP conceived the idea for the project and developed the approach, with contributions from TK. XP developed the computational methods and performed the simulations. TK provided guidance, mentorship and resources. XP and TK wrote the manuscript.

## Supporting information

**S1 Table. Ligand type frequencies in the binding site library.**

**S1 File. Summary tables of matching results to all fold families.**

## References

1. Feldmeier K, Hocker B. Computational protein design of ligand binding and catalysis. Curr Opin Chem Biol. 2013;17(6):929–33.

2. Feng J, Jester BW, Tinberg CE, Mandell DJ, Antunes MS, Chari R, et al. A general strategy to construct small molecule biosensors in eukaryotes. eLife. 2015;4.

3. Glasgow AA, Huang YM, Mandell DJ, Thompson M, Ritterson R, Loshbaugh AL, et al. Computational design of a modular protein sense-response system. Science (New York, NY. 2019;366(6468):1024–8.

4. Yang W, Lai L. Computational design of ligand-binding proteins. Current opinion in structural biology. 2017;45:67–73.

5. Zanghellini A, Jiang L, Wollacott AM, Cheng G, Meiler J, Althoff EA, et al. New algorithms and an in silico benchmark for computational enzyme design. Protein Sci. 2006;15(12):2785–94.

6. Jiang L, Althoff EA, Clemente FR, Doyle L, Rothlisberger D, Zanghellini A, et al. De novo computational design of retro-aldol enzymes. Science (New York, NY. 2008;319(5868):1387–91.

7. Rothlisberger D, Khersonsky O, Wollacott AM, Jiang L, DeChancie J, Betker J, et al. Kemp elimination catalysts by computational enzyme design. Nature. 2008;453(7192):190–5.

8. Tinberg CE, Khare SD, Dou J, Doyle L, Nelson JW, Schena A, et al. Computational design of ligand-binding proteins with high affinity and selectivity. Nature. 2013;501(7466):212–6.

9. Polizzi NF, Wu Y, Lemmin T, Maxwell AM, Zhang SQ, Rawson J, et al. De novo design of a hyperstable non-natural protein-ligand complex with sub-A accuracy. Nat Chem. 2017;9(12):1157–64.

10. Bick MJ, Greisen PJ, Morey KJ, Antunes MS, La D, Sankaran B, et al. Computational design of environmental sensors for the potent opioid fentanyl. eLife. 2017;6.

11. Dou J, Vorobieva AA, Sheffler W, Doyle LA, Park H, Bick MJ, et al. De novo design of a fluorescence-activating beta-barrel. Nature. 2018;561(7724):485–91.

12. Lucas JE, Kortemme T. New computational protein design methods for de novo small molecule binding sites. PLoS computational biology. 2020;16(10):e1008178.

13. Polizzi NF, DeGrado WF. A defined structural unit enables de novo design of small-molecule-binding proteins. Science (New York, NY. 2020;369(6508):1227–33.

14. Orengo CA, Pearl FM, Bray JE, Todd AE, Martin AC, Lo Conte L, et al. The CATH Database provides insights into protein structure/function relationships. Nucleic acids research. 1999;27(1):275–9.

15. Pan X, Thompson MC, Zhang Y, Liu L, Fraser JS, Kelly MJS, et al. Expanding the space of protein geometries by computational design of de novo fold families. Science (New York, NY. 2020;369(6507):1132–6.

16. Basanta B, Bick MJ, Bera AK, Norn C, Chow CM, Carter LP, et al. An enumerative algorithm for de novo design of proteins with diverse pocket structures. Proceedings of the National Academy of Sciences of the United States of America. 2020;117(36):22135–45.

17. Eswar N, John B, Mirkovic N, Fiser A, Ilyin VA, Pieper U, et al. Tools for comparative protein structure modeling and analysis. Nucleic acids research. 2003;31(13):3375–80.

18. Sillitoe I, Dawson N, Lewis TE, Das S, Lees JG, Ashford P, et al. CATH: expanding the horizons of structure-based functional annotations for genome sequences. Nucleic acids research. 2019;47(D1):D280–D4.

19. Leman JK, Weitzner BD, Lewis SM, Adolf-Bryfogle J, Alam N, Alford RF, et al. Macromolecular modeling and design in Rosetta: recent methods and frameworks. Nat Methods. 2020;17(7):665–80.

20. Huang PS, Boyken SE, Baker D. The coming of age of de novo protein design. Nature. 2016;537(7620):320–7.

21. Skolnick J, Gao M. Interplay of physics and evolution in the likely origin of protein biochemical function. Proceedings of the National Academy of Sciences of the United States of America. 2013;110(23):9344–9.

22. Skolnick J, Gao M, Zhou H. How special is the biochemical function of native proteins?. F1000Res. 2016;5.

23. Baker D. What has de novo protein design taught us about protein folding and biophysics? Protein Sci. 2019;28(4):678–83.

24. Davey JA, Damry AM, Euler CK, Goto NK, Chica RA. Prediction of Stable Globular Proteins Using Negative Design with Non-native Backbone Ensembles. Structure. 2015;23(11):2011–21.

25. Chaudhury S, Lyskov S, Gray JJ. PyRosetta: a script-based interface for implementing molecular modeling algorithms using Rosetta. Bioinformatics. 2010;26(5):689–91.

26. Park H, Bradley P, Greisen P, Jr., Liu Y, Mulligan VK, Kim DE, et al. Simultaneous Optimization of Biomolecular Energy Functions on Features from Small Molecules and Macromolecules. J Chem Theory Comput. 2016;12(12):6201–12.

27. Alford RF, Leaver-Fay A, Jeliazkov JR, O’Meara MJ, DiMaio FP, Park H, et al. The Rosetta all-atom energy function for macromolecular modeling and design. J Chem Theory Comput. 2017.

28. O’Boyle NM, Banck M, James CA, Morley C, Vandermeersch T, Hutchison GR. Open Babel: An open chemical toolbox. J Cheminform. 2011;3:33.

